# The effect of bioRxiv preprints on citations and altmetrics

**DOI:** 10.1101/673665

**Authors:** Nicholas Fraser, Fakhri Momeni, Philipp Mayr, Isabella Peters

## Abstract

A potential motivation for scientists to deposit their scientific work as preprints is to enhance its citation or social impact, an effect which has been empirically observed for preprints in physics, astronomy and mathematics deposited to arXiv. In this study we assessed the citation and altmetric advantage of bioRxiv, a preprint server for the biological sciences. We retrieved metadata of all bioRxiv preprints deposited between November 2013 and December 2017, and matched them to articles that were subsequently published in peer-reviewed journals. Citation data from Scopus and altmetric data from Altmetric.com were used to compare citation and online sharing behaviour of bioRxiv preprints, their related journal articles, and non-deposited articles published in the same journals. We found that bioRxiv-deposited journal articles received a sizeable citation and altmetric advantage over non-deposited articles. Regression analysis reveals that this advantage is not explained by multiple explanatory variables related to the article and its authorship. bioRxiv preprints themselves are being directly cited in journal articles, regardless of whether the preprint has been subsequently published in a journal. bioRxiv preprints are also shared widely on Twitter and in blogs, but remain relatively scarce in mainstream media and Wikipedia articles, in comparison to peer-reviewed journal articles.

## 2. Introduction

Preprints, typically defined as versions of scientific articles that have not yet been formally accepted for publication in a peer-reviewed journal, are an important feature of modern scholarly communication (Berg et al., 2016). Major motivations for the scholarly community to adopt the use of preprints have been proposed as early discovery (manuscripts are available to the scientific community earlier, bypassing the time-consuming peer review process), open access (manuscripts are publicly available without having to pay expensive fees or subscriptions) and early feedback (authors can receive immediate feedback from the scientific community to include in revised versions) (Maggio et al., 2018). An additional incentive for scholars to deposit preprints may be to increase citation counts and/or altmetric indicators such as shares on social media platforms. For example, recent surveys conducted by the Association for Computational Linguistics (ACL) and Special Interest Group on Information Retrieval (SIGIR), which investigated community members behaviours and opinions surrounding preprints, found that 32 and 15 % of respondents were respectively motivated to deposit preprints “to maximize the paper’s citation count” (Foster et al., 2017; Kelly, 2018).

A body of evidence has emerged which supports the notion of a citation differential between journal articles that were previously deposited as preprints and those that were not, with several studies concluding that arXiv-deposited articles subsequently received more citations than non-deposited articles in the same journals (Davis and Fromerth, 2007, Moed, 2007; Gentil-Beccot et al., 2010; Larivière et al., 2014). Multiple factors have been proposed as drivers of this citation advantage, including increased readership due to wider accessibility (the “open access effect”), earlier accumulation of citations due to the earlier availability of articles to be read and cited (the “early access effect”), authors preferential deposition of their highest quality articles as preprints (the “self-selection effect”), or a combination thereof (Kurtz et al., 2005). Whilst a citation advantage has been well documented for articles deposited to arXiv, the long-established nature of depositing preprints in physics, astronomy and mathematics may make it unsuitable to extend the conclusions of these studies to other subject-specific preprint repositories, where preprint deposition is a less established practice.

bioRxiv is a preprint repository aimed at researchers in the biological sciences, launched in November 2013 and hosted by the Cold Spring Harbor Laboratory (https://www.biorxiv.org/). As a relatively new service, it presents an interesting target for analysing impact metrics in a community where preprints have been less widely utilised in comparison to the fields of physics, astronomy and mathematics (Ginsparg, 2016). A recent study by Serghiou and Ioannidis (2018) provided initial insights into the potential citation and altmetric advantage of bioRxiv-deposited articles over non-deposited articles, finding that bioRxiv-deposited articles had significantly higher citation counts and altmetric scores than non-deposited articles.

In this study, we investigate citation and altmetric behaviour of bioRxiv preprints and their respective published papers, and compare them to papers not deposited to bioRxiv to determine if a citation and/or altmetric advantage exists. Our study builds on the study of Serghiou and Ioannidis (2018) in several ways: (1) we take into account a longer time period of analysis, from November 2013 to December 2017 (approximately one year longer than that analysed by Serghiou and Ioannidis (2018)), (2) we investigate longitudinal trends in citation behavior for preprints and their published papers, (3) we include a wider range of altmetric indicators including tweets, blogs, mainstream media articles, Wikipedia mentions and Mendeley reads, (4) we conduct regression analysis to investigate the influence of multiple factors related to publication venue and authorship, such as the journal impact factor, or number of co-authors per paper, which may have an effect on citation and altmetric differentials between articles deposited to bioRxiv and those not. Whilst we do not claim *causative* relationships in this study, we aim to shed light on factors that should be considered in discussions centered on preprint citation and altmetric advantages, and put our findings into the context of previous studies conducted on other preprint repositories.

## 3. Methods

### 3.1 Preprint and Article Metadata

Basic metadata of all preprints submitted to bioRxiv between November 2013 and December 2017 were harvested in April 2019 via the Crossref public Application Programming Interface (API) (N = 18,841), using the *rcrossref* package for R (Chamberlain et al., 2019). Links to articles subsequently published in peer-reviewed journals were discovered via three independent methods:

1. Via the ‘relationship’ property stored on the Crossref preprint metadata record. These links are maintained and routinely updated by bioRxiv through monitoring of databases such as Crossref and PubMed, or through information provided directly by the authors (personal correspondence with bioRxiv representative, October 2018). Each DOI contained in the ‘relationship’ property was queried via the Crossref API to retrieve the metadata record of the published article.
2. Via the publication notices published directly on the bioRxiv website (see, for example, https://doi.org/10.1101/248278). bioRxiv web pages were crawled in April 2019 using the *RSelenium* and *rvest* packages for R (Wickham, 2016; Harrison, 2019) and DOIs of published articles were extracted from the relevant HTML node of the publication notices.
3. Via matching of preprints records in Scopus (leveraging the data infrastructure of the German Competence Centre for Bibliometrics: http://www.forschungsinfo.de/Bibliometrie/en/index.php). Our matching procedure relied on direct correspondence of the surname and first letter of the given name of the first author, and fuzzy matching of the article title or first 100 characters of the abstract between the bioRxiv preprint and Scopus record. Fuzzy matching was conducted with the R package *stringdist* (van der Loo, 2018), using the Jaro distance algorithm and a similarity measure of 80 %. Matches were further validated by comparison of the author count of the preprint and Scopus record.

Overlapping links produced by the three separate methodologies (Figure 1) were merged to create a single set of preprint-published article DOI links. In rare cases of disagreement between methodologies (e.g. where the published paper DOI identified via the bioRxiv website differed to that identified via Crossref or our Scopus fuzzy-matching methodology), we prioritised the record from the bioRxiv website, followed by the Crossref record, with our Scopus fuzzy-matching methodology as the lowest priority. We discovered a small number of cases where authors had created separate records for multiple preprint versions rather than uploading a new version on the same record (e.g. https://doi.org/10.1101/122580 and https://doi.org/10.1101/125765). For these cases we selected the earlier posted record and discarded the later record from our dataset, to ensure that only a single non-duplicated published article exists for each preprint. Following these steps we produced a set of 12,767 links between deposited preprints and published articles, representing 67.8 % of all preprints deposited over the same time period.

**Figure 1:**
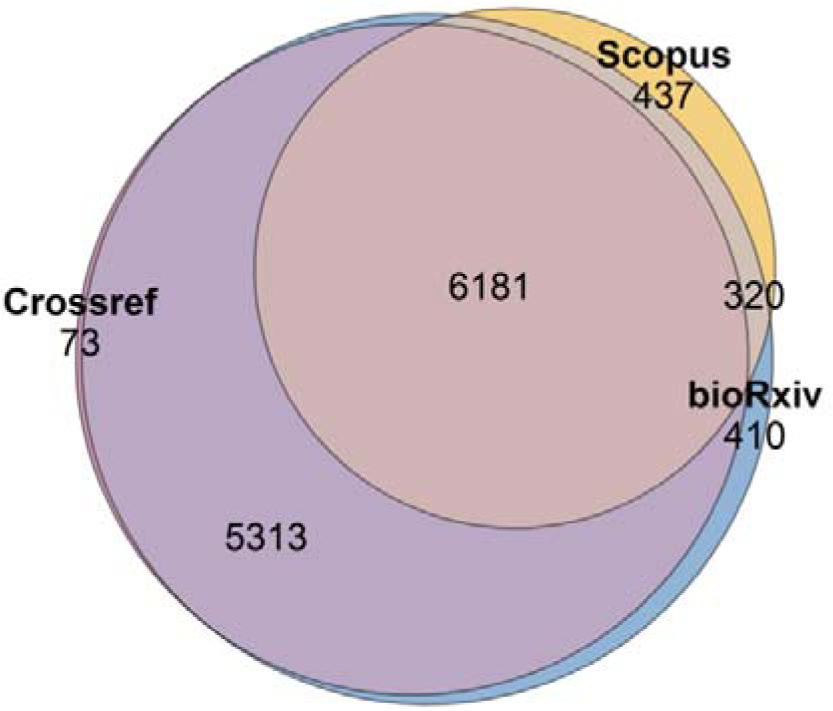
Proportional Venn diagram showing overlap between preprint-published article links discovered via three separate methodologies. ‘Crossref’ refers to those discovered via the Crossref ‘relationship’ property, ‘bioRxiv’ to those discovered via the bioRxiv website, and ‘Scopus’ to those discovered via fuzzy matching of preprint titles and abstracts to Scopus records.

### 3.2 Citation and altmetric analysis dataset

For the purposes of citation and altmetric analysis, we limited the set of journal articles retrieved in the previous step to those that were published in the 50-month period between November 2013 (coinciding with the launch of bioRxiv) and December 2017. We selected this time period as we use an archived Scopus database ‘snapshot’, which only partially covers articles published in 2018 (thus we only use years with full coverage). We further restricted the set of journal articles to those that could be matched to a record in Scopus via direct, case-insensitive correspondence between DOIs, to ‘journal’ publication types, ‘article’ or ‘review’ document types, and to articles with reference counts greater than zero, to reduce the rare incidence of editorial material incorrectly classified in Scopus as ‘article’ type documents.

Subsequently we built a control group of non-deposited articles for conducting comparative analysis. The control group was generated as follows: for each individual article within our bioRxiv-deposited group, we sampled a single random, non-deposited article published in the same journal and same calendar month. Articles in the control group were limited to ‘journal’ publication types, ‘article’ or ‘review’ document types, and records with reference counts greater than zero. We therefore generated a control group that matches our bioRxiv-deposited group in terms of journals and article ages.

A potential weakness of this matching procedure lies in the inclusion of articles published within large multidisciplinary journals (e.g. PLOS One, Scientific Reports), as it would be unwise to match a biology-focused article with an article from another discipline with drastically different publication and citing behaviours. For articles published in multidisciplinary journals, we therefore conducted an additional procedure prior to sampling, in which articles in both the bioRxiv-deposited and control groups were re-classified into Scopus subject categories based on the most frequently cited subject categories amongst their references (modified from the multidisciplinary article classification procedure used in Piwowar et al., 2018). Where categories were cited equally frequently, articles were assigned to multiple categories. For each bioRxiv-deposited article, a single random non-deposited article was sampled from the same journal-month and categories in the control group.

Following these steps, we produced an analysis dataset consisting of 7,087 bioRxiv-deposited and 7,087 non-deposited control articles.

### 3.3 Publication Dates

A methodological consideration when analysing citation data is in the treatment of publication dates. Publication dates for individual articles are reported by multiple outlets (e.g. by Crossref, Scopus and the publishers themselves), but often represent different publication points, such as the date of DOI registration, the Scopus indexing date, or the online and print publication dates reported by the publisher (Haustein et al., 2015). In our study, we implement the Crossref ‘created-date’ property as the canonical date of publication for all articles and citing articles in our datasets, in line with the approach of Fang and Costas (2018). The ‘created-date’ is the date upon which the DOI is first registered and can thus be considered a good proxy for the first online availability of an article at the publisher website. An advantage of this method is that we can report citation counts at a monthly resolution, as advocated by Donner (2018), which may be more suitable than reporting annual citation counts due to the relatively short time-span of our analysis period and rapid growth of bioRxiv. Created-dates of all preprints, articles and citing articles referenced in this study were extracted via the Crossref public API.

### 3.4 Citation Data

Metadata of citing articles were retrieved from Scopus for all articles in our bioRxiv-deposited and control groups. Citing articles were limited to those published over the time period of our analysis, November 2013 to December 2017. For each published article, we extract all citing articles and retrieve their Crossref created-date, to allow us to aggregate monthly citation counts. A consequence of this approach is that the maximum citation period of an article is variable, limited by the length of time between its publication, and the end of our analysis period in December 2017. For instance, an article published in December 2014 would have a maximum citation period of 36 months (from December 2014 to December 2017), whilst an article published in June 2017 would have a maximum citation period of 6 just months.

We additionally extracted records of articles directly citing preprints. Since preprints are not themselves indexed in Scopus, we utilised the Scopus raw reference data, which includes a ‘SOURCETITLE’ field including the location of the cited object. We queried the SOURCETITLE for entries containing the string ‘biorxiv’ (case-insensitive, partial matches), and retrieved a total of 4,826 references together with metadata of their Scopus-indexed citing articles. References were matched to preprints via fuzzy-matching of titles and/or direct matching of DOIs, although DOIs were only provided in a minority of cases. In total 4,387 references (90.9 %) could be matched to a bioRxiv preprint.

### 3.5 Altmetrics Data

Altmetric data, including tweets, blogs, mainstream media articles, Wikipedia, and Mendeley reads were retrieved for all deposited and non-deposited articles, as well as for preprints themselves, by querying their DOIs against the Altmetric.com API (https://api.altmetric.com/). Where no altmetric information was found for each indicator, counts were recorded as zero. Coverage amongst altmetric indicators was highest for Mendeley reads and Tweets, with 92 % and 91 % of published articles in our dataset receiving at least a single Mendeley read or Tweet. Coverage of Wikipedia mentions was lowest, with only 4 % of our articles being mentioned in Wikipedia.

### 3.6 Regression analysis

To investigate the influence of additional factors on a citation or altmetric differential between bioRxiv-deposited and non-deposited papers, we conducted regression analysis on citation and altmetric count data with a set of explanatory variables related to the article and its authorship. These variables include the journal impact factor (IF), article open access (OA) status, article type, first and last author country, first and last author academic age, and first and last author gender.

IF was calculated independently from Scopus citation data, following the formula:

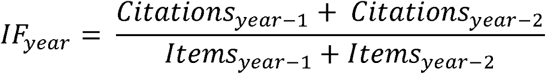

Note that items were limited to article and review document types, i.e. not including editorial material. Calculating IF independently ensures greater coverage of journals within our dataset compared to using the more commonly-known Journal Citation Reports produced by Clarivate Analytics. A manual comparison between the two datasets, however, suggests good agreement between the two methodologies.

Article OA status was determined by querying article DOIs against the Unpaywall API (https://unpaywall.org). Unpaywall is a service which locates openly available versions of scientific articles, via harvesting of data from journals and OA repositories. They provide a free API which can be queried via a DOI, returning a response containing information relating to the OA status, license and location of the OA article. We use the Boolean *‘is_oa’* resource returned by the Unpaywall API, which classifies articles as OA when the published article is openly available in any form, either on the publishers’ website or via an alternative repository (i.e. we do not distinguish between the ‘Gold’, ‘Green’ and ‘Hybrid’ routes of OA).

The country of the first and last author of each article was extracted from Scopus based upon the country in which the authors’ institution/s is/are based. For regression analysis, we classified authors into two categories: those having a US-based affiliation, and those not, following similar approaches employed by Gargouri et al. (2010) and Davis et al. (2008). Such an approach may not capture all of the fine-grain relationships between author countries and citations/altmetrics, however, it is notable that bioRxiv-deposited articles are generally over-represented by US-based authors: approximately 49 % of first and last authors of bioRxiv-deposited articles in our dataset had a US-based affiliation, whilst only around 38 % of first and last authors of non-deposited articles had a US-based affiliation.

The academic age of the first and last author of an article, used as a proxy for academic seniority, was determined from the difference between the publication year of the paper in question, and the year of the authors’ first recorded publication in Scopus. Whilst there are limitations to this approach, for example we may not detect authors who publish preferentially in edited volumes not indexed in Scopus, the year of first publication has be found to be a good predictor for both the academic and biological age of a researcher in multiple subject areas (Nane, Larivière and Costas, 2017). To obtain the year of the first recorded publication, we retrieved authors’ publication histories using the Scopus author ID, an identifier assigned automatically by Scopus to associate authors with their publication *oeuvres*. The author ID aims to disambiguate authors based upon affiliations, publication histories, subject areas and co-authorships (Moed et al., 2013). The algorithm aims at higher precision than recall; that is to say, articles grouped under the same author ID are likely to belong to a single author, but the articles of an author may be split between multiple author IDs.

Author gender was inferred using the web service Gender API (https://gender-api.com). Author first names were extracted from Scopus and stripped of any leading or trailing initials (e.g. “Andrea B.” would become “Andrea”). Gender API predicts gender using a database of >2 million name-gender relationships retrieved from governmental records and data crawled from social networks (Santamaria and Mihaljevic, 2018). The service accepts parameters for localization, which we included from our previously defined dataset of author countries. Gender assignments are returned as “male”, “female”, or “unknown”. Where localized queries returned “unknown”, we repeated the query without the country parameter. For our data, we were able to assign genders to 94.4 % of first authors, and 94.8 % of last authors.

Regression analysis was conducted for citation counts (for 6-month, 12-month, and 24-month citation windows) and altmetric counts (for tweets, blogs, mainstream media articles, Wikipedia mentions, and Mendeley reads) using a negative binomial regression model, with the full set of explanatory variables as described above. A negative binomial regression model is more suitable for over-dispersed count data (as is the case with citation and altmetric count data) than a linear regression model (Ajiferuke and Famoye, 2015). Regression was conducted using the R package *MASS* (Venables and Ripley, 2002).

## 4. Results and Discussion

### 4.1 bioRxiv submissions and publication outcomes

Depositions of preprints to bioRxiv grew exponentially between November 2013 and December 2017 (Figure 2). Of the 18,841 preprints posted between 2013 and 2017, our matching methodology identified 12,767 preprints (67.7 %) that were subsequently published in peer reviewed journals. This is a slightly higher rate than the 64.0 % reported by Abdill and Blekhman (2019), which may be due to our analysis occurring later (thus allowing more time for preprints to be published), as well as our more expansive matching methodology which did not rely solely on publication notices on the bioRxiv websites. These results from bioRxiv are broadly similar to those of Larivière et al. (2014) in the context of ArXiv, who found that 73 % of ArXiv preprints were subsequently published in peer-reviewed journals, with the proportion decaying in more recent years as a result of the delay between posting preprints and publication in a journal. The stability of the proportion of bioRxiv preprints that proceeded to journal publication between 2013 and 2016 additionally suggests that the rapid increase in the number of preprint submissions was not accompanied by any major decrease in the quality (or at least, the ‘publishability’) of preprints over this time period.

**Figure 2:**
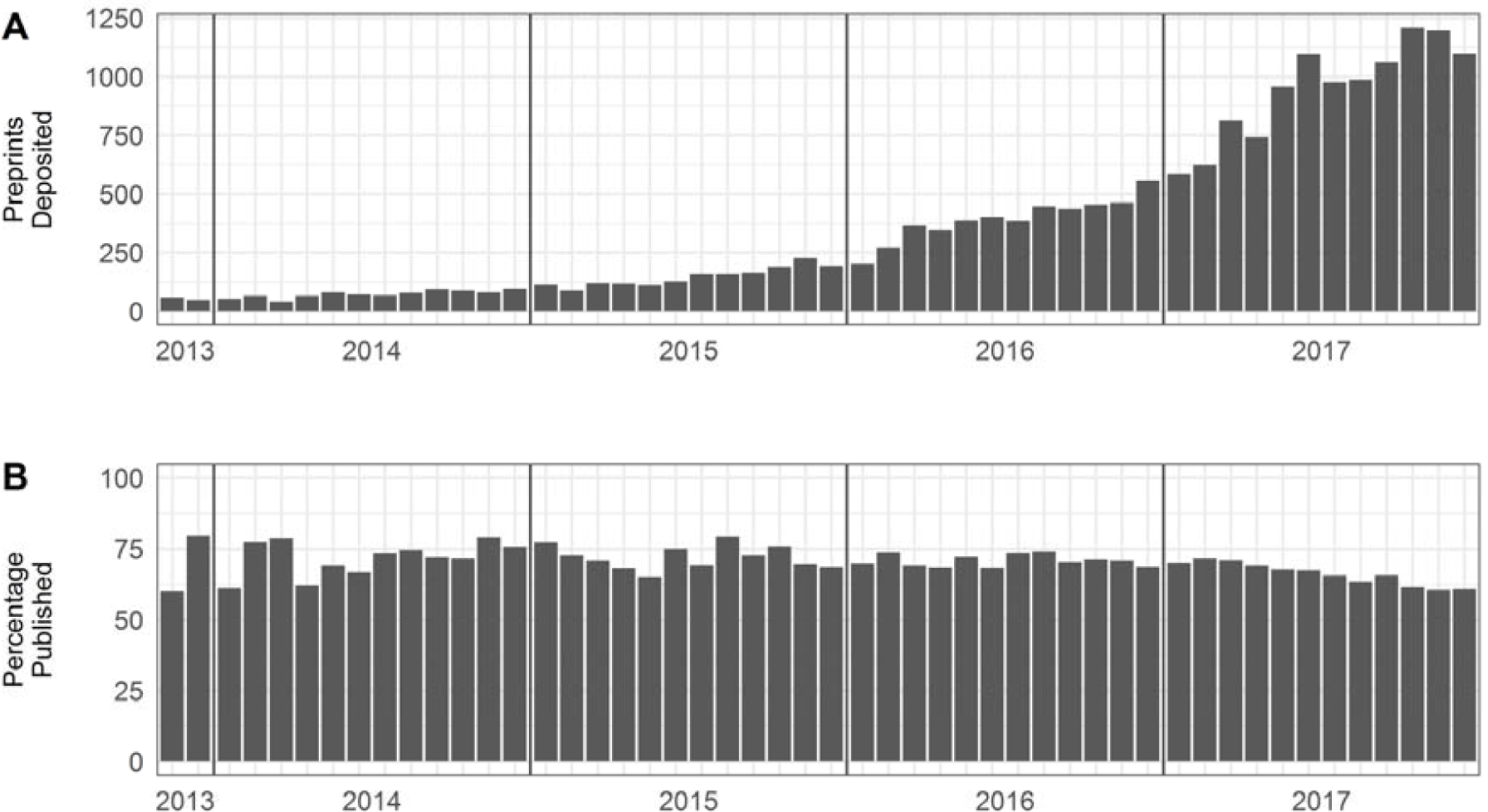
Development of bioRxiv submissions and publication outcomes over time. **(A) S**ubmissions of preprints to bioRxiv. **(B)** Percentage of bioRxiv preprints subsequently published in peer-reviewed journals.

The median delay time between submission of a preprint and publication was found to be 154 days, in comparison to the 166 days reported by Abdill and Blekhman (2019) – the difference can likely be explained by the different points of publication used – whilst we used only the Crossref ‘created-date’, Abdill and Blekhman (2019) prioritised the ‘published-online’ date, and the ‘published-print’ date when ‘published-online’ was not available, only using the ‘created-date’ as a final option. It should be noted that neither of these calculated delay times is representative of the average *review* time of a manuscript submitted to a journal, as authors may not submit their manuscript to a journal immediately on deposition of a preprint, and manuscripts may be subject to several rounds of rejection and resubmission before publication. Nonetheless, the delay time calculated by both our approach and that of Abdill and Blekhman (2019) reveals that preprints effectively shorten the time to public dissemination of an article by 5-6 months compared to the traditional journal publication route.

### 4.2 Citations Analysis

#### 4.2.1 bioRxiv citation advantage

For the time period November 2013 to December 2017, we retrieved a total of 47,169 citations to journal articles that were previously deposited to bioRxiv, versus 29,298 citations to articles in our non-deposited control group. These numbers give a crude citation advantage of bioRxiv-deposited articles of 61.0 % over non-deposited articles published in the same journal and month. A similar crude citation advantage of bioRxiv-deposited articles was also reported by Serghiou and Ioannidis (2018), despite the usage of different citation data sources - in our study we use citation data derived from Scopus whilst Serghiou and Ioannidis (2018) use citation data derived from Crossref. A recent analysis has found similar overall coverage of publications and citations of both Scopus and Crossref (Harzing, 2019), however there are still some major gaps in Crossref citation coverage due to non-support of certain Crossref members of the Open Citations Initiative (https://i4oc.org), most notably Elsevier, which may still introduce systematic bias into large-scale citation analyses.

To explore how the bioRxiv citation advantage develops over time following publication, we compared average monthly citations per paper (*Cpp*) for each group for the 36 months following journal publication (Figure 3). Citation counts were aggregated at a monthly level for each article, and then counts were log-transformed to normalize the data and reduce the influence of papers with high citation counts (following Thelwall (2016) and Ruocco et al. (2017)). *Cpp* was calculated by taking the mean of the log-transformed citation counts of all articles within a group:

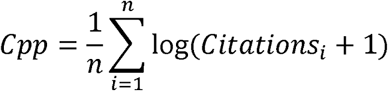

We limited our citation window to 36 months due to the small number of articles that were published sufficiently early in our analysis to allow longer citation windows. In general terms, we observe an acceleration of the citation rates of both groups within the first 18 months following publications, and an approximate plateau in citation rates between 18 and 36 months. However, the results demonstrate a clear divergence between the two groups beginning directly at the point of publication; at 6 months post-publication the *Cpp* of bioRxiv-deposited articles is 29 % higher than the non-deposited articles, with the monthly advantage growing to 40 % by 12 months post-publication. Between 18-36 months, the citation differential stabilises, with the *Cpp* of the bioRxiv-deposited group remaining ∼50 % higher than the control group.

**Figure 3:**
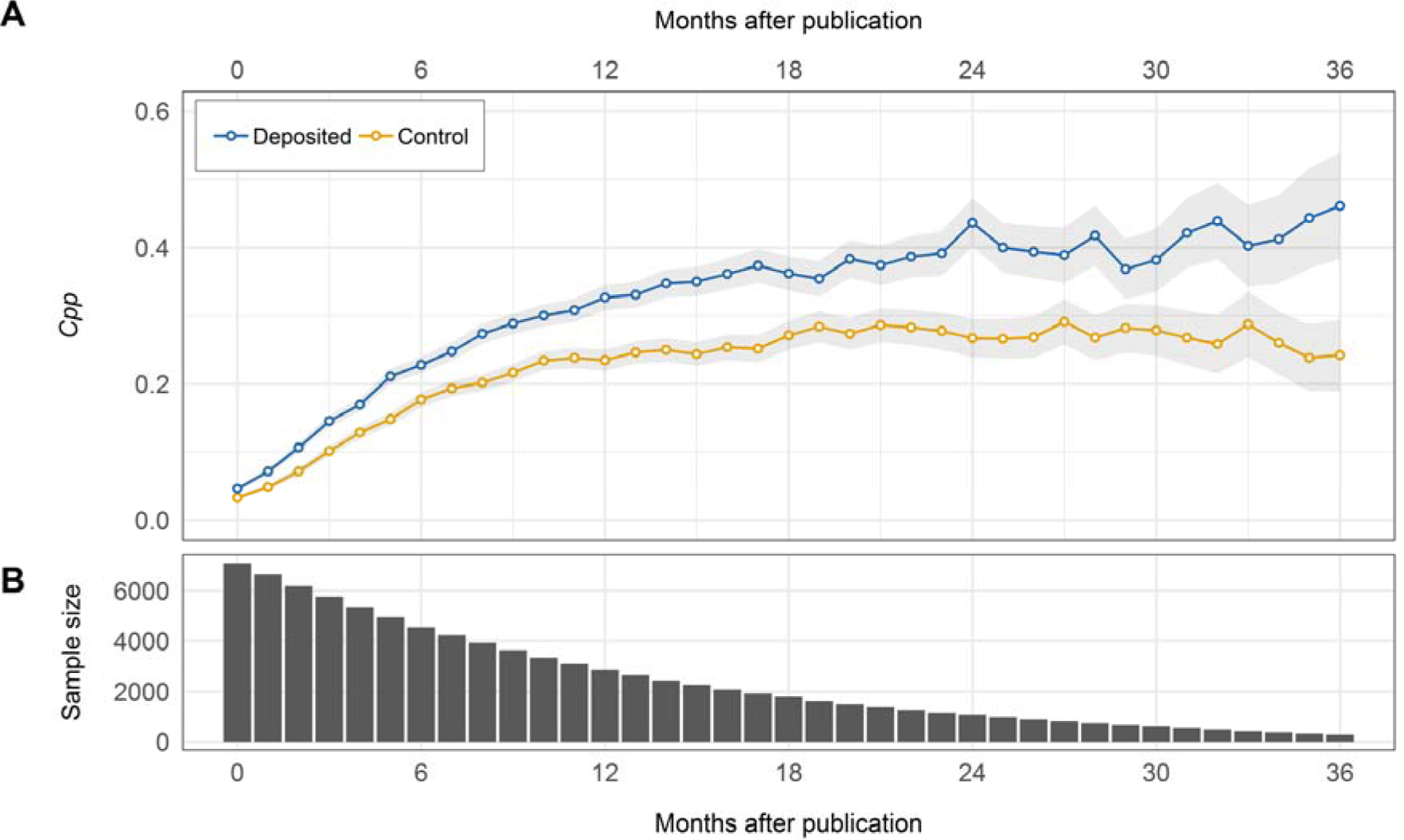
Monthly citation rates of bioRxiv-deposited and non-deposited control articles. **(A)** Calculated *Cpp* of bioRxiv-deposited articles (blue line) and non-deposited control articles (yellow line) as a function of months following publication. Grey shading represents 95 % confidence intervals. **(B)** Sample size of each group at each respective time interval. Sample sizes are equal for both groups.

The stability of the citation differential between bioRxiv-deposited and non-deposited articles after 18 months means that we cannot attribute the citation advantage solely to an *early access* effect, where articles with preprints receive a short-term acceleration in citations due to their earlier availability and thus longer period to be read and cited. If this were the case we would expect citation rates of both groups to converge after a period of time, as was reported by Moed (2007) in the context of preprints deposited to ArXiv’s Condensed Matter section. In the Moed (2007) study, monthly average citation rates of ArXiv-deposited and non-deposited articles converged after approximately 24 months, whilst our data show no sign of similar behaviour. Conversely, other studies tracking longitudinal changes in citation rates of articles deposited in other arXiv communities have found less support for an early access effect (Henneken et al., 2006; Gentil Beccot et al., 2009), with citations for deposited articles remaining higher than non-deposited articles for >5 years following publication.

An alternative explanation for the citation advantage of bioRxiv-deposited articles is that of a *quality* effect, which can be manifested either as a quality bias driven by users self-selecting their highest quality articles to deposit (Kurtz et al., 2005; Davis and Fromerth, 2007), or as a quality advantage where high quality articles which are more likely to be selectively cited anyway are made more accessible, thus further boosting their citedness (Gargouri et al., 2010).

We test for a quality advantage through a secondary analysis in which articles were divided into categories on the basis of their respective journal impact factors (IF). Whilst it is well recognised that the IF is not a good measure of the quality of an individual article (Cagan, 2013), it remains an important predictor of academic job success in biomedicine (van Dijk, Manor, Carey; 2014), and can thus be considered as a proxy for researchers’ *perception* of the highest quality outlets to submit their work, i.e. an author is more likely to submit their perceived higher quality work to a high-IF journal. Each article was assigned an IF on the basis of its journal and year of publication, and then articles divided into quartiles on the basis of the IF of the journal in which it was published. The upper quartile (‘High IF’) contained articles in journals with IFs above 7.08, the second highest quartile (“Med-High IF”) between 7.08 and 4.53, the second lowest quartile (“Med-Low IF”) between 4.53 and 3.33, and the lower quartile (“Low IF”) lower than 3.33. Monthly citation rates were calculated as previously, within each IF quartile (Figure 4). We observe that the *absolute* citation advantage grows faster in the High IF group, but the *relative* advantage remains remarkably consistent between all four groups, particularly during the first 18 months following publication. If the citation advantage was driven primarily by the articles perceived as researchers highest quality articles, we would expect to see a citation advantage manifested primarily amongst articles in high IF journals, with relatively equal citation rates between bioRxiv-deposited and non-deposited articles in low IF journals. Our data, however, do not appear to support this view.

**Figure 4:**
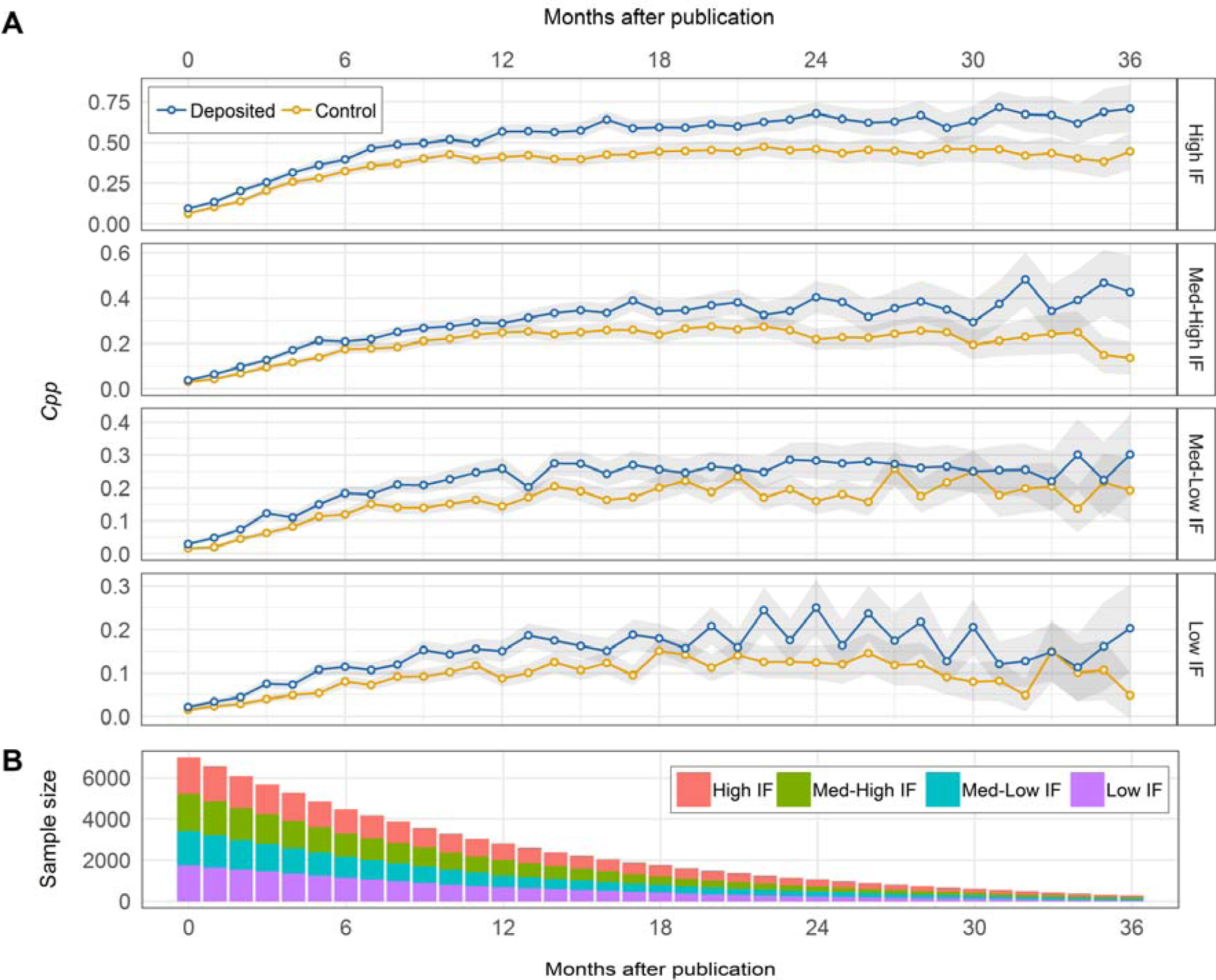
Monthly citation rates of bioRxiv-deposited and non-deposited control articles grouped by IF quartiles. High IF articles are classified as those in a journal with an IF >7.08, Med-High IF between 7.08-4.53, Med-Low IF between 4.53-3.33, and low IF <3.33. **(A)** Calculated *Cpp* of bioRxiv-deposited articles (blue line) and non-deposited control articles (yellow line) as a function of months following publication for High IF (upper panel), Med-High IF (second top panel), Med-Low IF (third top panel), and Low UF (lower panel) journals. Grey shading represents 95 % confidence intervals. **(B)** Sample size of each group at each respective time interval. Sample sizes and IF distributions are equal for the bioRxiv-deposited and control groups.

#### 4.2.2 Citations to preprints

In addition to retrieving citations to journal articles, we also retrieved details of 4,387 citations made directly to preprints themselves. Of these, 2,107 citations were made to preprints that were subsequently published as journal articles, whilst the remaining 2,280 citations were made to preprints that remain unpublished. Figure 5 shows a comparison between the *Cpp* of preprints that have subsequently been published in journals, and those that remain unpublished, for a 24 month citation window following deposition of the preprint. Citations to preprints that have been published increase sharply in the first 6 months following deposition, and thereafter decrease, likely a result of other authors preferentially citing the journal version of an article over the preprint. Similar findings have been reported for ArXiv preprints (Brown, 2001; Henneken, 2007; Larivière, 2014). It is interesting to note that in the early months following deposition, unpublished preprints are not cited any less than their published counterparts, and continued to accrue citations many months after deposition, even in the absence of an accompanying journal article. Citing of unpublished preprints is in itself a relatively new development in biological sciences; the National Institutes of Health (NIH), for example, only adopted a policy allowing scientists to cite preprints in grant applications in March 2017 (https://grants.nih.gov/grants/guide/notice-files/NOT-OD-17-050.html), and some journals have only recently allowed authors to cite preprints directly in their reference lists (see, e.g. Stoddard and Fox (2019)). Although the number of citations to bioRxiv preprints is still dwarfed by those to journal articles (the *Cpp* of preprints is more than an order of magnitude less than the *Cpp* of the respective publisher papers), the growing willingness of authors to cite unreviewed preprints may factor into ongoing debates surrounding the role of peer review and maintaining the integrity of scientific reporting.

**Figure 5:**
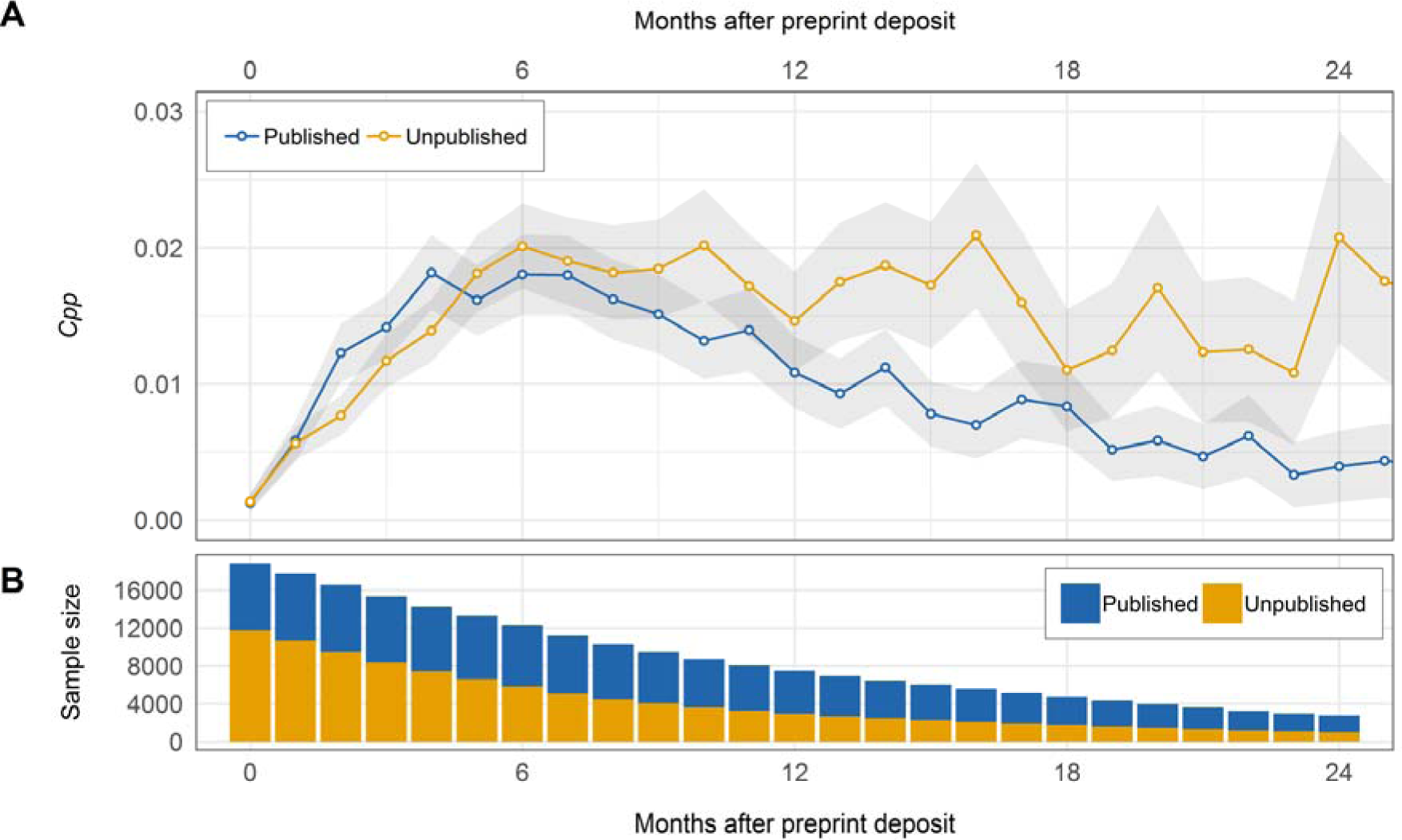
Monthly citation rates of bioRxiv preprints. Preprints are divided into two categories: those which have subsequently been published in peer-reviewed journals, and those which remain unpublished. **(A)** Calculated *Cpp* of published (blue line) and unpublished (yellow line) bioRxiv preprints as a function of months following preprint deposition. Grey shading represents 95 % confidence intervals. **(B)** Sample sizes at each respective time interval.

Figure 6 shows the distribution of monthly citation rates to preprints as a function of time before and after the publication of the journal article, i.e. negative citation months indicate the preprint was cited before the journal article was published, and vice versa. Citations appear to become more frequent in the months shortly preceding publication of the journal article, and fall sharply thereafter. A small number of preprints continue to accrue citations more than two years after publication of the journal article, although the origin of these citations is not clear: they may be citations from authors who do not have access to journal publications requiring subscriptions, from authors who remain unaware that a preprint has been published elsewhere or authors failing to update their reference management software with the record from the journal article. A similar analysis of citation aging characteristics of arXiv preprints found that citations to preprints decay rapidly following publication of the journal article (Larivière et al., 2014), whilst reads of arXiv preprints through the NASA Astrophysics Data System also dropped to close to zero following publication of the peer-reviewed article, attributed to authors preferring to read the journal article over the preprint (Henneken et al., 2007).

**Figure 6:**
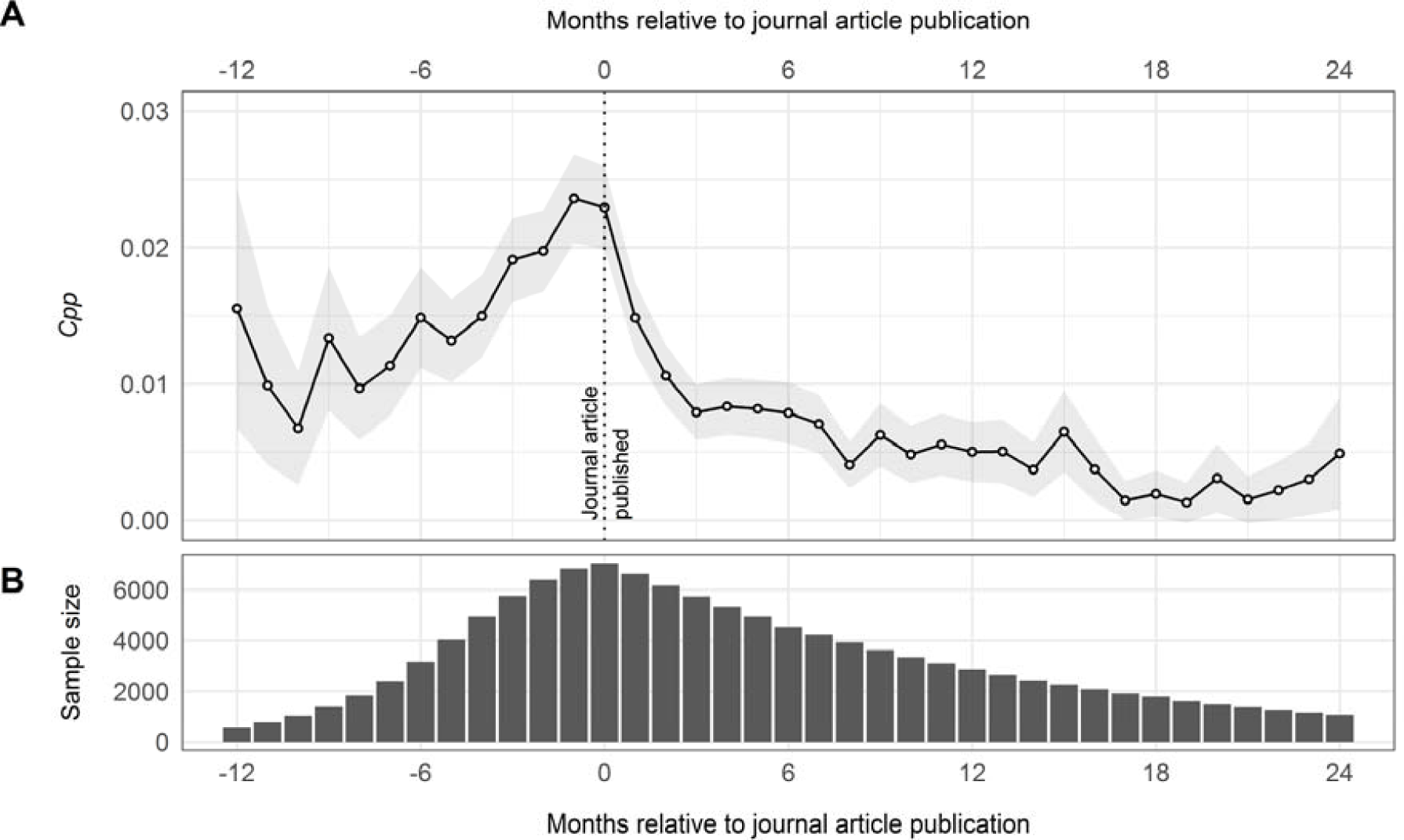
Monthly citation rates of preprints before and after journal publication. **(A)** Calculated *Cpp* of bioRxiv preprints for the 12 months prior to, and 24 months following journal publication. Grey shading represents 95 % confidence interval. **(B)** Sample size of preprints at each time interval.

### 4.3 Altmetrics

Altmetric data were retrieved from Altmetric.com and aggregated for all bioRxiv-preprints, bioRxiv-deposited articles and non-deposited control articles (Figure 7). Since altmetrics accrue rapidly in comparison to citations (Bornmann, 2014), we do not aggregate altmetrics into time windows as is more common with citation analysis. Coverage of altmetrics (i.e. the proportion of articles that received at least one count in the various altmetric sources) for bioRxiv-preprints, bioRxiv-deposited articles and non-deposited control articles were 99.7, 96.3 and 87.4 %, respectively. It should be noted that the high coverage of altmetrics in bioRxiv-preprints is in large part due to the automatic tweeting of newly published bioRxiv-preprints by the official bioRxiv twitter account (https://twitter.com/biorxivpreprint), although we cannot discount automatic tweeting by publishers, journals or individuals for the other categories.

**Figure 7:**
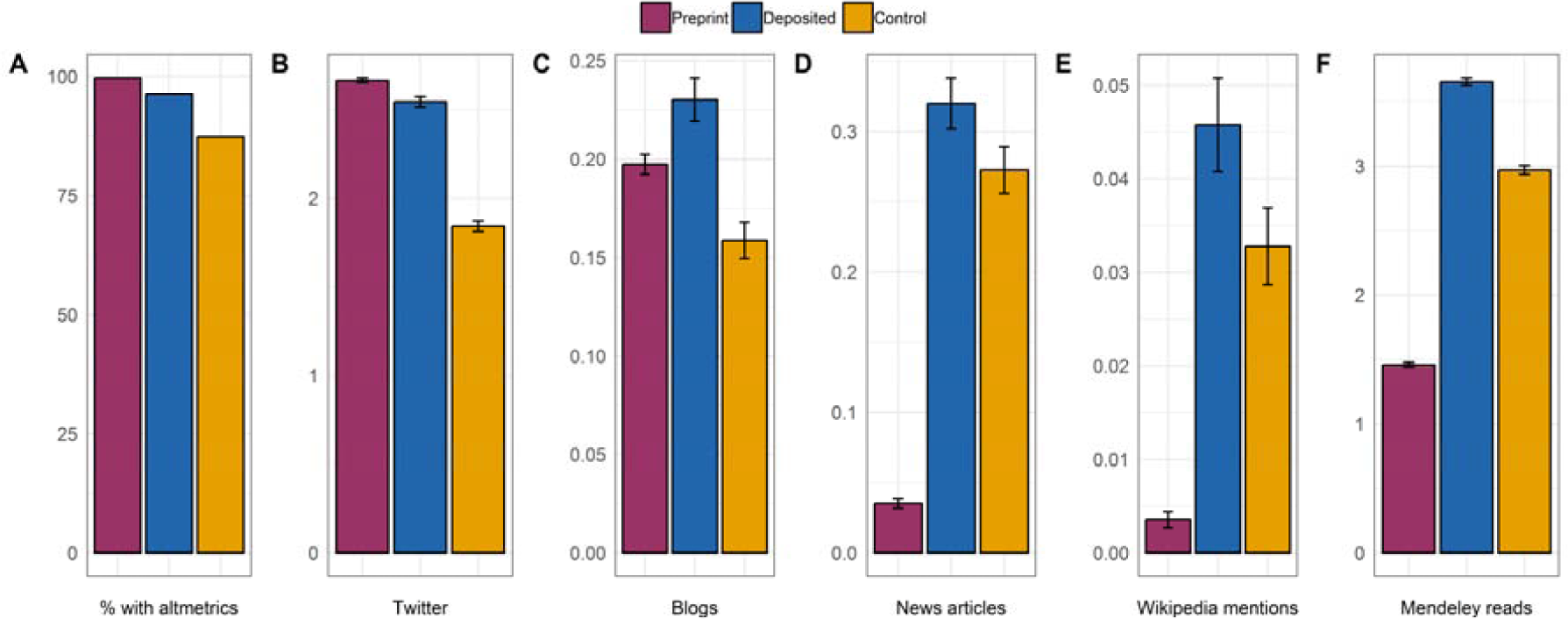
Altmetric coverage and counts of bioRxiv-preprints, bioRxiv-deposited articles and non-deposited control articles. Altmetric counts were log-transformed prior to reporting. **(A)** Percentage of articles associated with any altmetric event covered by Altmetric.com. **(B)** Mean count of tweets. **(C)** Mean count of blog mentions. **(D)** Mean count of mentions in mainstream media articles. **(E)** Mean count of Wikipedia mentions. **(F)** Mean count of Mendeley reads.

Figure 7B-F show mean (log-transformed) counts of tweets, blogs, mainstream media articles, Wikipedia mentions and Mendeley reads, for bioRxiv-preprints, bioRxiv-deposited articles and non-deposited control articles. For all of these data sources, mean counts were higher for the bioRxiv-deposited articles than the non-deposited control articles, indicating that articles that have previously been shared as a preprint are subsequently shared more in various online platforms, in agreement with the previous results of Serghiou and Ioannidis (2018). Mean counts of tweets and blog mentions were broadly similar in bioRxiv-preprints and bioRxiv-deposited articles, but strikingly lower in mainstream news articles and mentions in Wikipedia. This may suggest that whilst bioRxiv preprints are widely shared in ‘informal’ social networks by colleagues and peers, they are currently less accepted in ‘formal’ public outlets where peer-reviewed articles remain the preferred source.

In a similar vein to our previous citation analysis, we conducted a secondary analysis for altmetrics by dividing articles into quartiles on the basis of their journal IF, and comparing altmetric coverage and counts for bioRxiv-deposited and non-deposited control articles (Figure 8). In all cases, altmetric coverage and mean altmetric counts remain higher for the bioRxiv-deposited articles, indicating a preference for sharing of articles previously deposited as preprints over those not deposited. Notable differences are observed between the IF categories, where High IF articles receive more altmetric attention in general that Low IF articles. A similar positive correlation between preprint downloads and IF was reported by Abdill and Blekhman (2019), although in the absence of a ubiquitous source of download data for journal articles, we cannot extend these findings to compare downloads between bioRxiv-deposited and non-deposited articles. As with our citation analysis, the *absolute* differences in altmetric counts between the bioRxiv-deposited and non-deposited articles vary greatly between IF categories, but the *relative* differences remain relatively similar, indicating that there is also no general quality effect driving the bioRxiv altmetric advantage.

**Figure 8:**
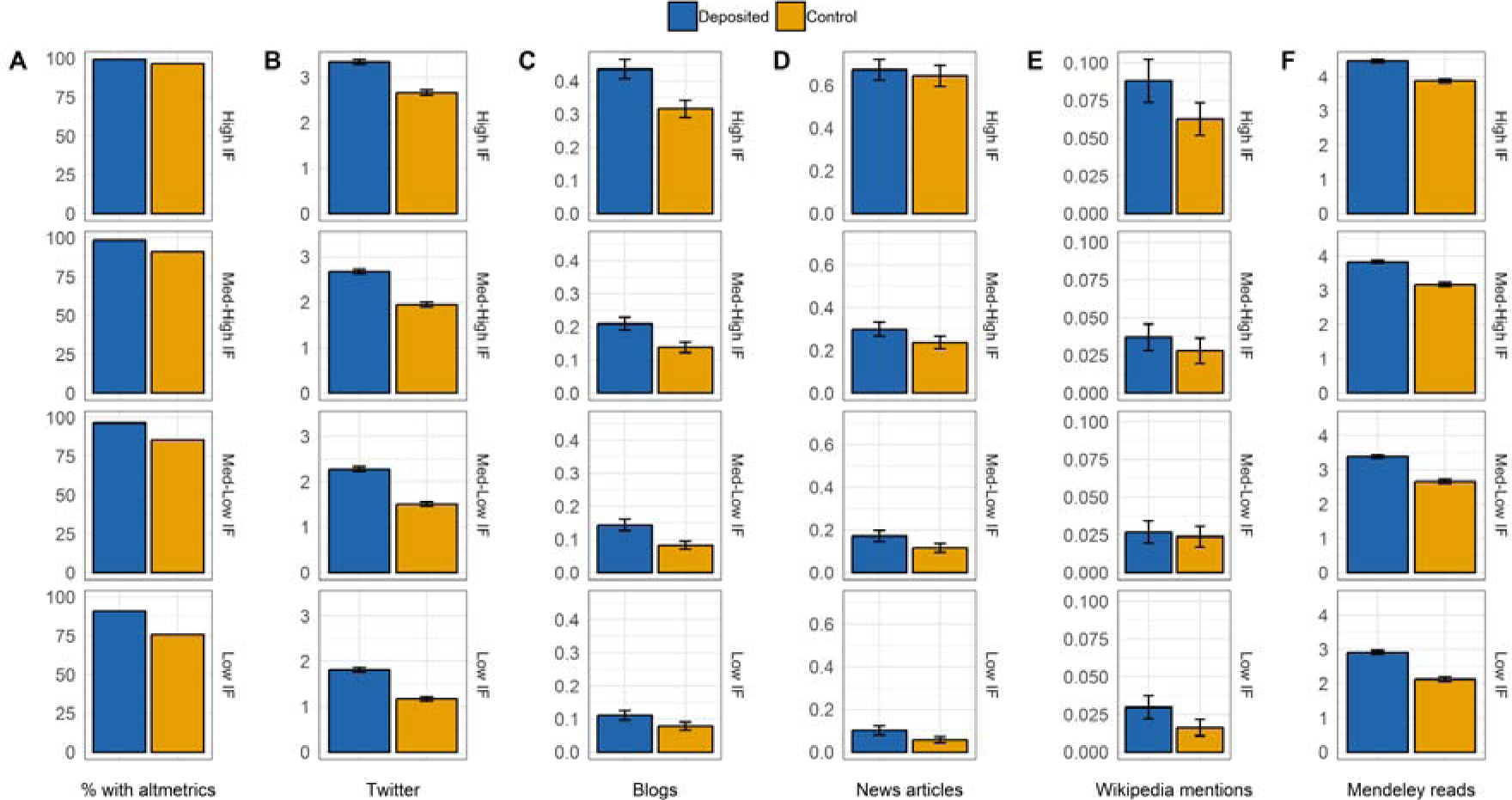
Altmetric coverage and counts of bioRxiv-deposited articles and non-deposited control articles grouped by IF quartiles. High IF articles are classified as those in a journal with an IF >7.08, Med-High IF between 7.08-4.53, Med-Low IF between 4.53-3.33, and low IF <3.33. Upper panels show results for High IF articles, second top panel for Med-High IF articles, third top panel for Med-Low IF articles, and the lower panel for Low IF articles. Altmetric counts were log-transformed prior to reporting. **(A)** Percentage of articles associated with any altmetric event covered by Altmetric.com. **(B)** Mean count of tweets. **(C)** Mean count of blog mentions. **(D)** Mean count of mentions in mainstream media articles. **(E)** Mean count of Wikipedia mentions. **(F)** Mean count of Mendeley reads.

### 4.4 Regression analysis

Results in the previous sections suggest a sizeable citation and altmetric advantage of depositing preprints to bioRxiv, in the absence of consideration of other factors related to publication venue and authorship. In a second step we therefore conducted regression analysis to determine the effect of bioRxiv deposition on citations and altmetrics when controlling for multiple explanatory variables (summarised in Table 1). These explanatory variables are not exhaustive, as citations and altmetrics can be influenced by a number of additional variables which we do not account for (Tahamtan et al., 2016; Didegah et al., 2018), and do not take into account certain *immeasurable* characteristics of an article such as its underlying quality or the quality of the authors themselves. Thus, we refrain from claiming a definitive causative relationship between bioRxiv deposition and a citation or altmetric advantage. However, these variables may help to shed some light on factors which influence citation or altmetric differentials, which may be considered and explored in future studies.

**Table 1:** Summary statistics for set of 10 explanatory variables included in our regression analysis.

| Characteristic | Variable | bioRxiv-deposited<br>(N = 7,087) | Control<br>(N = 7,087) |
| --- | --- | --- | --- |
| **Journal Impact Factor** | Median Journal Impact Factor | 4.53 | 4.53 |
| **OA article** | % papers that are OA | 88.1 | 83.5 |
| **Review article** | % review articles | 2.79 | 5.32 |
| **Author count** | Median author count per article | 5 | 6 |
| **First author from USA** | % US first authors | 49.3 | 37.4 |
| **Last author from USA** | % US last authors | 49.5 | 37.6 |
| **First author academic age** | Median first author academic age (years) | 5 | 5 |
| **Last author academic age** | Median last author academic age (years) | 17 | 19 |
| **First author gender** | % articles with female first authors | 29.9 | 36.0 |
| **Last author gender** | % articles with female last authors | 18.0 | 23.9 |

Summary statistics for explanatory variables (Table 1) reveal some key differences between articles that were deposited to bioRxiv, and those that were not. Articles deposited to bioRxiv are more likely to subsequently be published under an OA license than non-deposited article. Here we used the most inclusive categorisation of OA provided by Unpaywall, and did not distinguish between types of OA such as Gold and Green OA. However, given that our two samples are matched with respect to journals, differences arising in OA coverage must result from author choices to make their paper open through Hybrid OA options in subscription journals, or through Green OA self-archiving (e.g. in institutional repositories).

We found that 2.8 % of bioRxiv-deposited articles were classified as ‘review’ type documents, despite the bioRxiv website stating that review and hypothesis articles should not be posted, and that “manuscripts that solely summarize existing knowledge or present narrative theories are inappropriate” (https://www.biorxiv.org/about/FAQ). In contrast, 5.3 % of articles in our non-deposited control group are review article types.

The median number of authors per paper is lower for articles deposited to bioRxiv than those not; this is a somewhat surprising, as it may be logically inferred that the more papers an author has, the more likely it is to be deposited as a preprint at the request/suggestion of one of the authors. For the first and last authors of an article, US authors were found to be overrepresented in the bioRxiv-deposited articles compared to non-deposited control articles, which may partly be a result of bioRxiv being a US-based platform, as well as institutional and/or funding policies in the US encouraging the deposition of preprints. Median academic age for both groups was found to be similar for first authors, but last authors were slightly younger in the bioRxiv-deposited group than the non-deposited group, indicating that preprints may be a phenomenon driven more by the younger generation of scientists. Female authors were found to be underrepresented compared to male authors for both groups, although the imbalance was greater in the bioRxiv-deposited group than the non-deposited group; of first authors in the bioRxiv-deposited group, only 29.9 % were female, falling to 18.0 % for last authors. The finding that female authors are underrepresented as authors in biomedical fields in general is in agreement with previous research (e.g. Larivière et al., 2013), however the mechanism by which female authors are even less well represented amongst preprint authors is not clear. Similar findings were reported from a survey of authors conducted by the Association for Computational Linguistics; whilst 31 % of total respondents were female, only 12.5 % of those who state they always or often post to preprint servers were female (Foster et al., 2017).

Model parameters for regression analysis on citation counts using 6, 12 and 24 month citation windows are summarised in Table 2. Note that for regression analysis on citation data, our sample sizes decreased as the citation window increased, due to the number of articles which had a sufficient citation length within our period of analysis. Where values were missing (e.g. when we were unable to determine the gender of an author), we removed both the bioRxiv-deposited and matched non-deposited articles from our analysis, to maintain balance of publication ages between groups. We interpret significance at the p<0.005 level, following the recommendations of Benjamin et al. (2018).

**Table 2:** Negative binomial regression output reporting effects of 11 explanatory variables on citation counts. Regression analysis was undertaken at 3 key time intervals: 6 months post publication, 12 months post publication and 24 months post publication. Values in bold indicate significance at the p<0.005 level, following the recommendations of Benjamin et al. (2018).

| Variable | 6-month citations (N = 7,654) |  |  | 12-month citations (N = 4,784) |  |  | 24-month citations (N = 1,838) |  |  |
| --- | --- | --- | --- | --- | --- | --- | --- | --- | --- |
| | B | $\beta$ | p | B | $\beta$ | p | B | $\beta$ | p |
| Intercept | **-1.193** | - | **<0.001** | -0.000 | - | 0.996 | **1.200** | - | **<0.001** |
| **Deposited to bioRxiv** | **0.421** | **0.049** | **<0.001** | **0.422** | **0.015** | **<0.001** | **0.529** | **0.007** | **<0.001** |
| IF | **0.086** | **0.110** | **<0.001** | **0.088** | **0.036** | **<0.001** | **0.080** | **0.013** | **<0.001** |
| OA | **0.376** | **0.029** | **<0.001** | **0.356** | **0.008** | **<0.001** | **0.297** | **0.002** | **<0.001** |
| Author Count | **0.014** | **0.043** | **<0.001** | **0.012** | **0.013** | **<0.001** | **0.006** | **0.002** | **<0.001** |
| Review Article | **0.264** | **0.012** | **<0.001** | **0.347** | **0.005** | **<0.001** | **0.342** | **0.002** | **0.001** |
| US First Author | 0.012 | 0.001 | 0.862 | 0.125 | 0.004 | 0.066 | **0.436** | **0.005** | **<0.001** |
| US Last Author | 0.092 | 0.011 | 0.172 | -0.038 | -0.001 | 0.573 | **-0.406** | **-0.005** | **<0.001** |
| First Author Academic Age | 0.003 | 0.005 | 0.190 | 0.001 | 0.001 | 0.588 | -0.006 | -0.001 | 0.148 |
| Last Author Academic Age | 0.006 | 0.008 | 0.035 | 0.006 | 0.002 | 0.044 | 0.008 | 0.001 | 0.110 |
| Female First Author | -0.082 | -0.08 | 0.016 | **-0.135** | **-0.004** | **<0.001** | **-0.200** | **-0.002** | **<0.001** |
| Female Last Author | 0.022 | 0.002 | 0.573 | -0.010 | 0.000 | 0.802 | -0.063 | -0.001 | 0.299 |

For all three citation windows, the bioRxiv-deposited status, IF, OA status, author count and review article status were found to be significant predictors of citations. For 12-month citations, the first author gender was also found to be a significant predictor, and for 24-month citations the country of the first and last author (US or not US) were additionally found to be significant predictors. First and last author academic age, and the gender of the last author did not significantly predict citation counts in any of the citation windows.

By taking the exponent of the regression coefficient, *Exp(B)*, we calculate an Incidence Rate Ratio (IRR) of bioRxiv deposition of 1.52 for a 6-month citation window. That is to say, when controlling for all other explanatory variables, bioRxiv-deposited articles receive a citation advantage of 52 % over non-deposited articles. The citation advantage increases to 53 % for a 12-month citation window, and to 70 % for a 24-month citation window. Our regression analysis showed however, that several other characteristics of articles not related to bioRxiv deposition status had large effects on citation rates, for example for each unit increase in journal IF, citations increased by ∼8-9 % in all citation windows, whilst an article being OA increased citations by ∼45 % for a 6 month citation window, although the effect appears to reduce with time, only increasing citations by and 35 % for a 24 month citation window. These results clearly demonstrate that any attempts to quantify a citation advantage of a single platform or repository such as bioRxiv need to carefully consider other factors influencing citation counts in their analyses.

Model parameters for regression analysis on altmetrics are summarised in Table 3. For all altmetric indicators investigated (tweets, blogs, mainstream media articles, Wikipedia mentions, Mendeley reads), bioRxiv deposition status, IF and number of authors were significant predictors of altmetric counts. OA status was additionally a significant predictor across all altmetric indicators with the exception of Wikipedia mentions. Calculated IRRs suggest that bioRxiv deposition has the largest impact on tweets – bioRxiv deposited articles received 93 % more tweets when controlling for the set of explanatory variables than non-deposited articles (47 % for blog feeds, 24 % for media mention, 36 % for Wikipedia mentions, 75 % for Mendeley reads). As with our citation analysis, we do not aim to establish a causative link between bioRxiv deposition and altmetric indicators, but nonetheless our results show that bioRxiv-deposited articles are shared significantly more in online communities than non-deposited articles, even when controlling for multiple factors related to the article and its authorship.

**Table 3:** Negative binomial regression output reporting effects of 11 explanatory variables on various altmetric indicators including tweets, blog feeds, media mentions, Wikipedia mentions and Mendeley reads. Values in bold indicate significance at the p<0.005 level, following the recommendations of Benjamin et al. (2018).

| Variable | Tweets (N = 12,054) |  |  | Blog Feeds (N = 12,054) |  |  | Media (N = 12,054) |  |  | Wikipedia (N = 12,054) |  |  | Mendeley (N = 12,054) |  |  |
| --- | --- | --- | --- | --- | --- | --- | --- | --- | --- | --- | --- | --- | --- | --- | --- |
| | B | $\beta$ | p | B | $\beta$ | p | B | $\beta$ | p | B | $\beta$ | p | B | $\beta$ | p |
| Intercept | 1.328 |  | <0.001 | -2.637 | - | <0.001 | -2.507 | - | <0.001 | -3.974 | - | <0.001 | 2.695 | - | <0.001 |
| Deposited to bioRxiv | 0.656 | 0.006 | <0.001 | 0.383 | 0.147 | <0.001 | 0.219 | 0.017 | 0.001 | 0.304 | 0.258 | 0.003 | 0.558 | 0.002 | <0.001 |
| IF | 0.103 | 0.010 | <0.001 | 0.096 | 0.394 | <0.001 | 0.143 | 0.115 | <0.001 | 0.076 | 0.691 | <0.001 | 0.101 | 0.004 | <0.001 |
| OA | 0.454 | 0.003 | <0.001 | 0.670 | 0.173 | <0.001 | 0.957 | 0.049 | <0.001 | 0.322 | 0.184 | 0.049 | 0.327 | 0.001 | <0.001 |
| Author Count | 0.005 | 0.001 | <0.001 | 0.007 | 0.072 | <0.001 | 0.029 | 0.058 | <0.001 | 0.013 | 0.285 | <0.001 | 0.003 | 0.000 | <0.001 |
| Review Article | 0.257 | 0.001 | <0.001 | -0.037 | -0.005 | 0.763 | -0.560 | -0.016 | 0.002 | 0.436 | 0.144 | 0.066 | 0.777 | 0.001 | <0.001 |
| US First Author | 0.052 | 0.000 | 0.316 | 0.247 | 0.094 | 0.016 | 0.425 | 0.032 | 0.005 | 0.183 | 0.154 | 0.408 | 0.146 | 0.001 | <0.001 |
| US Last Author | 0.096 | 0.001 | 0.063 | 0.099 | 0.038 | 0.331 | 0.021 | 0.002 | 0.891 | -0.231 | -0.195 | 0.298 | -0.008 | 0.000 | 0.848 |
| First Author Academic Age | 0.024 | 0.002 | <0.001 | 0.015 | 0.0563 | <0.001 | 0.011 | 0.010 | 0.066 | 0.030 | 0.286 | <0.001 | 0.004 | 0.000 | 0.011 |
| Last Author Academic Age | -0.008 | -0.001 | <0.001 | -0.013 | -0.057 | 0.001 | -0.003 | -0.003 | 0.599 | 0.004 | 0.036 | 0.677 | -0.008 | 0.000 | <0.001 |
| Female First Author | -0.038 | -0.000 | 0.116 | -0.015 | -0.005 | 0.767 | 0.237 | 0.017 | 0.001 | -0.210 | -0.167 | 0.059 | -0.113 | 0.000 | <0.001 |
| Female Last Author | -0.031 | -0.000 | 0.264 | -0.096 | -0.030 | 0.101 | 0.085 | 0.005 | 0.308 | -0.065 | -0.045 | 0.609 | -0.070 | 0.000 | 0.003 |

Our regression results reveal several other interesting differences in the behaviour of altmetric indicators for articles in biological sciences. For example, review articles are significantly and positively correlated with numbers of tweets and Mendeley reads, but significantly and negatively correlated with the number of media mentions. This may show that articles reviewing and summarising previous knowledge are highly shared amongst networks of academics, they are not deemed particularly ‘newsworthy’ in comparison to more original research. With respect to author academic ages, both first author academic ages are found to positively predict mentions in tweets and blog feeds, but conversely the last author academic ages negatively predict mentions and tweets. Gender is found to have no significant effect on tweets, blog posts and Wikipedia mentions, but the gender of the first author is a positive predictor of news mention (articles with a female first author receive 27 % more mentions in mainstream media), whilst the gender of the first and last author negatively predict Mendeley reads. These are just a few examples of factors which show that individual altmetric indicators represent activity of different online communities (a full investigation of which is outside the scope of this study; see Haustein et al. (2015) and Didegah et al. (2018) for further discussion) and should thus be considered in isolation (instead of, e.g., aggregated Altmetric.com scores) in future studies attempting to understand the relationship between altmetrics and preprint deposition behaviour.

## 5. Conclusions

We have found empirical evidence that journal articles which have previously been posted as a preprint on bioRxiv receive more citations and more online attention than articles published in the same journals which were not deposited, even when controlling for multiple explanatory variables. In terms of citations, the advantage is immediate and long-lasting – even after three years following publication, bioRxiv-deposited articles continue to accrue citations at a higher rate than non-deposited articles. Our finding of a preprint citation advantage is in agreement with previous research conducted on arXiv, suggesting that there may be a general advantage of depositing preprints not limited to a single long-established repository. More research is needed to establish the exact cause of the citation and altmetric advantage. However, our results do not implicate a clear early access or quality effect in driving this advantage, which may point to access itself being the driver. Further research should dive deeper into understanding motivations of researchers to deposit their articles to bioRxiv, for example through qualitative survey and interviews, which will shed light on factors related to author bias and self-selection of articles to deposit.

We additionally investigated longitudinal trends in citation behaviour of preprints themselves, finding that preprints are being directly cited regardless of whether they have been published in a peer-reviewed journal or not, although there is a strong preference to cite the published article over the preprint when it exists. Preprints are also shared widely on Twitter and on blogs, in contrast to mainstream media articles and Wikipedia where published journal articles still dominate, suggesting that there remains some reluctance to promote un-reviewed research to public audiences. In the continuing online debates surrounding the value of preprints and their role in modern scientific workflows, our results provide support for depositing preprints as a means to extend the reach and impact of work in the scientific community. This may help to motivate and encourage authors, some of whom remain sceptical of preprint servers, to publish their work earlier in the research cycle.

## 6. Acknowledgements

This work is supported by BMBF project OASE, grant number 01PU17005A. Note that a shortened ‘work in progress’ version of this work, entitled “Examining the citation and altmetric advantage of biorXiv preprints”, was submitted as a conference paper to be presented at the 17^th^ International Conference on Scientometrics and Informatics (ISSI 2019; Rome, September 2-5, 2019).

## 7. Author Contributions

Conception and design: NF, FM, PM, IP. Acquisition of data: NF, FM. Analysis and interpretation of data: NF, FM, PM, IP. Drafting and revising of article: NF, FM, PM, IP.

## 8. Data Availability

Data and code used for the analysis presented in this paper can be found at https://github.com/nicholasmfraser/biorxiv.

